# Is HDL-c plasma concentration a possible marker of HIV replication? A cross-sectional analysis in untreated HIV-infected individuals accessing HIV care in Italy

**DOI:** 10.1101/2023.06.23.546265

**Authors:** Stefania Piconi, Martina Bottanelli, Giulia Marchetti, Andrea Gori, Antonella Castagna, Nicola Squillce, Stefania Cicalini, Giancarlo Orofino, Francesca Ceccherini-Silverstein, Antonio Di Biagio, Antonella d’Arminio Monforte, Alessandro Cozzi-Lepri, Icona Foundation Study Cohort

## Abstract

**Aims:** HIV infection is associated with dyslipidemia and an increased risk for cardiovascular diseases. HIV Nef protein downregulates the generation of nascent HDL. The interplay between HIV-RNA, HDL-c level and CD4/CD8 ratio in naïve HIV patients remains to be elucidated.

**Methods:** We included untreated persons living with HIV (PLWH) of the ICONA Foundation Study cohort if they also had ≥2 viral load (VL) measurements prior to ART initiation. We performed unadjusted correlation and linear regression analyses evaluating the effect of VLset on HDL-C. Vlset and CD4/CD8 ratio were fit in the log_10_ scale, while HDL-c, was fitted in the untransformed raw scale.

**Results:** We included 3,980 untreated PLWH. Fifty-eighty (1.5%) were aviremic. We observed a negative correlation between HDL-c and VLset (Pearson R^2^=0.03), from fitting an unadjusted linear regression model -8.5 mg/dl (95% CI: -15,9 --0,84 p<0.03). There was a dose-response relationship between HDL-c levels and VLset, however, this association was somewhat attenuated after further controlling for gender. Despite a positive correlation between HDL-c and CD4/CD8 ratio, the HDL-c plasma concentration does not satisfy the criteria for a strong surrogate marker.

**Conclusions:** Our data show that HDL-c plasma concentration is significantly lower per higher level of VLSet although this was in part explained by gender. Further analyses should be promoted to better understand the molecular mechanisms that underline the relationship between HIV replication, HDL-c formation, and diseases progression.

## Introduction

HIV infection has been associated with changes in lipid concentration, characterized by decreased levels of high-density lipoprotein cholesterol (HDL-c) and increased levels of low-density lipoprotein cholesterol (LDL-c), total cholesterol (TC) and triglycerides (TGL) [1-3].

Antiretroviral therapy (ART) increases HDL-c concentration by eliminating active viral replication, although this association might be confounded by demographic factors [4-6]. In addition, ART use leads to a rise in TC and LDL-c that typically exceeds pre-infection levels, whereas the recovery of HDL-c may be incomplete [2]. HDL-c is considered protective against the development of atherosclerosis because it removes atherogenic lipid molecules from foam-cells to the liver, facilitating its elimination in the intestinal tract [reverse cholesterol transport (RCT)], and it has also several antioxidant and anti-inflammatory properties which can help prevent LDL-c oxidation and inflammatory cell migration [7]. Consequently, ART and non-ART related lipid alteration, associated with chronic inflammation and adipose tissue dysfunction, can be clearly considered as one of the possible explanations for the increased risk of cardiovascular disease (CVD) events reported for people living with HIV (PLWH), compared to uninfected controls [8-12].

In addition, there are several immunological mechanisms trough which HDL-c has shown to have a protective role, particularly in sepsis, due to its critical intermediary step in lipid-based pathogen clearance, bacterial toxin binding and disposal [13-16], monocyte activation, macrophage and dendritic-cell migration, release of inflammatory cytokines [17-18] and inhibition of vascular and intercellular adhesion molecule expression [19].

Actually, it is known that cholesterol is a key component of cell membrane and virus envelope, and cholesterol-rich microdomains, known as lipid rafts, on host cell plasma membranes have an important role in viral entry and budding: in fact, it was demonstrated that cholesterol-depleting molecules, such as methyl-B-cyclodextrin, inhibit the cellular entry of several viruses, such as HIV-1, rotaviruses and coronaviruses [20]. Focusing on lipid alteration during HIV infection, the change in HDL-c would suggest that there are several steps of HIV replication that critically depend on cholesterol metabolism. The molecular confirmation of this hypothesis is offered by Mujavar’s [21] in vitro results. According to these results, the Nef HIV protein impairs ATP-binding cassette transporter A1 (ABCA-1) dependent cholesterol efflux from human macrophages generating several consequences, such as: cholesterol accumulation within monocytes (foam-cells transformation), reduction of HDL-c plasma concentration, increased virus budding (due to the rise of cytoplasmatic lipid rafts) and lastly an increase in HIV replication. Later in vitro studies with LXR-α agonists (TO-901317), a strong stimulator of ABCA-1 expression, showed an improvement of cholesterol efflux from HIV-infected T lymphocytes and macrophages associated with a reduction of HIV replication in both cell types. The effect of this antagonist is remarkably reduced in ABCA-1 defective T-cells of a patient with Tangier disease [22]. Furthermore, HIV ΔNef infection in vivo resulted in much lower VL and in a milder presentation of several elements of immunological dysfunction compared to patients infected with WT HIV [23]. Lipidomics techniques have also allowed the characterization of the lipidome of enveloped viruses. By this way, HIV lipid envelope has been observed to be different from the producer cell plasma membrane, suggesting that viruses bud from specialized membrane subdomains, which are enriched in particular lipids [24].

The evidence summarised above, supports the notion that plasmatic HDL-c is a should be biochemical marker which is likely to be related to HIV viral budding and inflammation. With this analysis, we aimed to corroborate, in the setting of real-life untreated HIV-infection, the association between VLset and lipids (such as total cholesterol and HDL-c plasma concentrations), and whether VLset mediated HDL-c changes might also correlate with immunological parameters of HIV progressions, such as CD4/CD8 ratio.

## Materials and Methods

### Study population

In this retrospective cross-sectional study, we included untreated HIV-infected people enrolled in the ICONA Foundation cohort. The main aim was to evaluate the association between HDL-c plasma concentration and VL set-point in absence of ART; a secondary objective was to evaluate the association between HDL-c levels and markers of HIV disease progression like CD4/CD8 ratio. We included people for whom ≥2 viral load (VL) measurements prior to ART initiation were available. The viral set point (VLset) was defined as the mean of the first two VL and the date of the 2^nd^ value chosen as the index date for this cross-sectional analysis. Participants with an estimated VLset <50 copies/mL were labelled as ‘aviremic’ and the remaining group as ‘viremic’. People who had started statin therapy prior to the index data and those without a value of HDL over 3 months of the index data were excluded. All laboratory markers test results were included in the correlation analyses if measured within 6 months of the index Vlset date.

### Ethical considerations

The Icona Foundation study was approved by the Ethics Committees (institutional review board) of each participating institution. All of the individuals enrolled provided a written informed consent at the time of the enrolment. All procedures of the study were performed in accordance with the 1964 Helsinki declaration and its later amendments.

### Statistical analyses

Characteristics of the study population were described overall and after stratification according to VLset (≤50 copies/mL vs. >50 copies/mL). The distribution of categorical factors was compared using a chi-square test and median values of continuous factors using the non-parametric Mann-Whitney test. Box plots were used to depict the full distribution (Q1, Q3, median, range) of lipid markers across the two groups. Unadjusted Pearson correlation coefficient was also used to test the hypothesis of a linear relationship between VLSet and lipids.

In order to control for potential confounding factors, a multivariable analysis was conducted for total cholesterol and HDL-c for which an univariable difference between groups was detected. In particular, the association between VLSet (main exposure) and HDL-c (primary outcome) and total cholesterol (secondary outcome) was evaluated by fitting a linear regression model after controlling for a minimal set confounders chosen *a priori* including gender, age, CD4/CD8 ratio, HCV status (detection of HCVAb), and AIDS diagnosis. Total cholesterol was essentially chosen as a negative control. This list of measured potential confounders was put together using both axiomatic knowledge and literature review. In order to further assess the robustness of the results against potential unmeasured confounding bias, the e-value was calculated and compared to the magnitude of the mean difference seen for predictors showing the strongest association with the outcome (25).

HDL-c and total cholesterol, which both showed a symmetric distribution, were fitted in the untransformed raw scale. VLSet instead was fitted in three ways: i) comparing people with ≤50 copies/mL (aviremic) vs. >50 copies/mL (viremic); ii) using the log_10_ scale and iii) after splitting the study population in groups using pre-specified HIV-RNA clinical cut-offs to evaluate a potential dose-response effect.

In addition, a refined model has been hypothesised for a third outcome: the CD4/CD8 ratio. In this model, on the basis of the results of the main analysis, BMI was the only confounder of the association between VLSet and CD4/CD8 ratio, while HDL-C was a mediator, i.e. some of the total effect of VLSet on CD4/CD8 is assumed to be explained by a variation in HDL-C. This was visually described using a direct acyclic graph (DAG, Figure 1). A mediation analysis was formally performed using the ‘medeff’ command in Stata 15. All other results were obtained from using SAS version 9.4 (Carey, USA).

**Figure 1.**
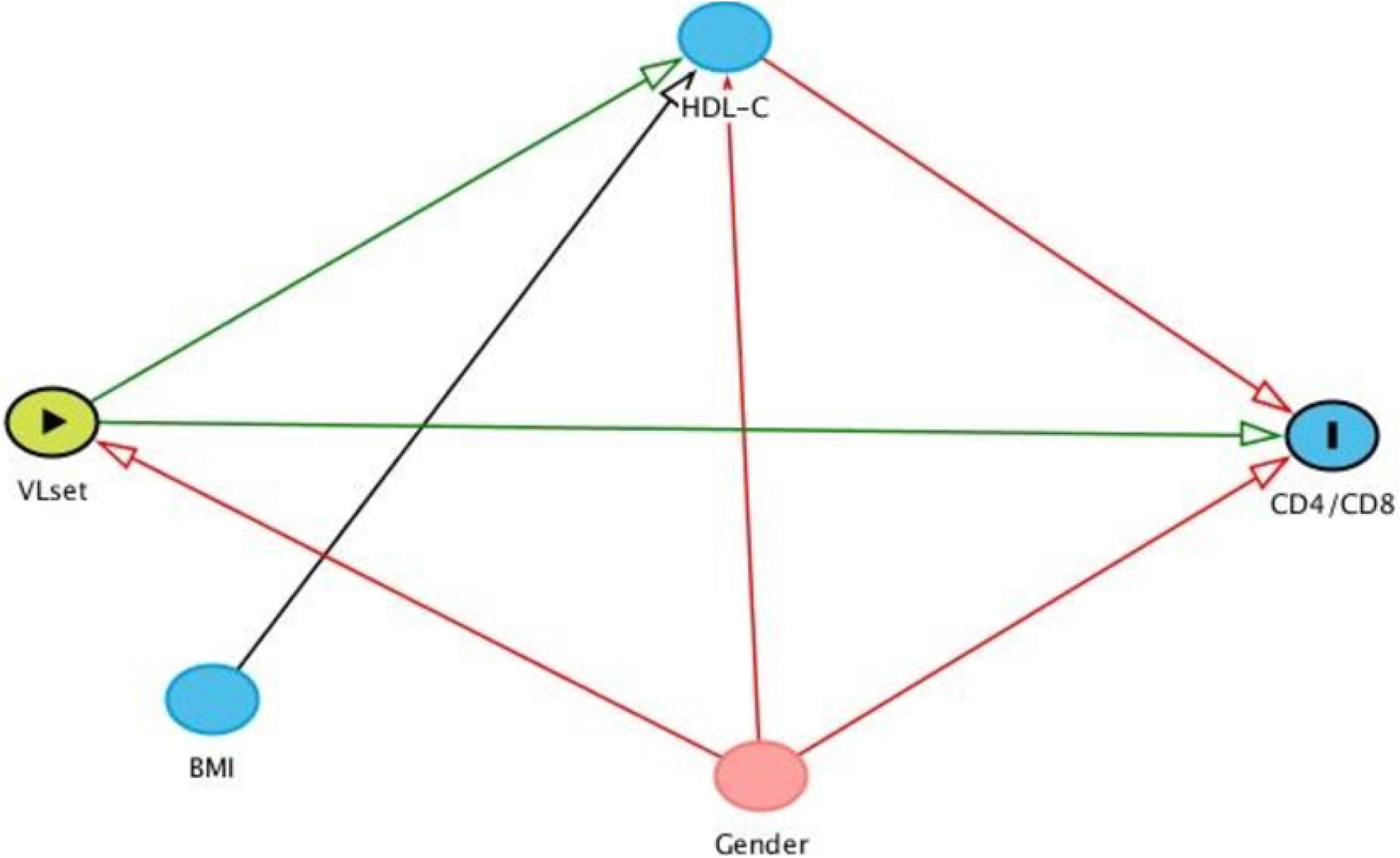
DAG of the mediation model with outcome CD4/CD8.

## Results

### Study population

The clinical and demographic characteristics of HIV positive patients enrolled in the study are shown in Table 1. We included 3,980 HIV ART-naive individuals, 58 patients (1.5%) spontaneously aviremic and 3,922 (98.5%) viremic patients, respectively. As shown in Table 1, the group of aviremic patients were significantly older [aviremic vs. viremic median 41 (IQR: 35, 48) vs. 37 (IQR: 30, 44) years p= 0.005] and with more females [aviremic vs. viremic 26 (45%) vs. 749 (19%) p<.001]. Furthermore, as expected, the aviremic patients presented higher TCD4 cell counts [aviremic vs. viremic 766 (IQR: 546, 1001) cells/mm^3^ vs. 535 (IQR: 384, 707) cells/mm^3^ p<0,001], and significantly lower TCD8 cells counts [aviremic vs. viremic 732 (IQR: 499, 997) cells/mm^3^ vs 984 (IQR: 718, 1352) cells/mm^3^ p<0,001] and VL [aviremic vs viremic 1.40 (IQR: 1.30, 1.66) log_10_ copies/ml vs 4.36 (IQR: 3.79, 4.85) log10 copies ml p<0.001] compared to viremic.

**Table 1.** Characteristics of the study populations stratified by HIV-RNA group.

| Characteristics | Viral load set point (copies/mL) |  |  |  |
| --- | --- | --- | --- | --- |
|  | 0-50<br>N= 58 | >50<br>N= 3922 | p-value* | Total<br>N= 3980 |
| **Gender, n (%)** |  |  | <.001 |  |
| Female | 26 (44.8%) | 749 (19.1%) |  | 775 (19.5%) |
| **Mode of HIV Transmission, n (%)** |  |  | <.001 |  |
| PWID | 14 (24.1%) | 356 (9.2%) |  | 370 (9.4%) |
| MSM | 12 (20.7%) | 1953 (50.2%) |  | 1965 (49.8%) |
| Heterosexual contacts | 28 (48.3%) | 1365 (34.8%) |  | 1393 (35.0%) |
| Other/Unknown | 4 (6.9%) | 214 (5.5%) |  | 218 (5.5%) |
| **Ethnicity, n (%)** |  |  | 0.002 |  |
| Caucasian | 44 (75.9%) | 3518 (89.7%) |  | 3562 (89.5%) |
| South America | 6 (10.3%) | 178 (4.5%) |  | 184 (4.6%) |
| Africa | 8 (13.8%) | 188 (4.8%) |  | 196 (4.9%) |
| Asian | 0 (0.0%) | 38 (1.0%) |  | 38 (1.0%) |
| **BMI** |  |  | 0.057 |  |
| Median (IQR) | 24 (22, 27) | 23 (21, 25) |  | 23 (21, 25) |
| **Smoking, n (%)** |  |  | 0.026 |  |
| No | 33 (56.9%) | 1590 (40.5%) |  | 1623 (40.8%) |
| Current | 20 (34.5%) | 1612 (41.1%) |  | 1632 (41.0%) |
| Unknown | 5 (8.6%) | 720 (18.4%) |  | 725 (18.2%) |
| **CNS diagnosis, n (%)** |  |  | 0.915 |  |
| Yes | 5 (8.6%) | 354 (9.0%) |  | 359 (9.0%) |
| **HBsAg, n (%)** |  |  | 0.427 |  |
| Negative | 39 (67.2%) | 2922 (74.5%) |  | 2961 (74.4%) |
| Positive | 0 (0.0%) | 5 (0.1%) |  | 5 (0.1%) |
| Not tested | 19 (32.8%) | 995 (25.4%) |  | 1014 (25.5%) |
| **HCVAb, n (%)** |  |  | 0.094 |  |
| Negative | 31 (53.4%) | 2618 (66.8%) |  | 2649 (66.6%) |
| Positive | 8 (13.8%) | 346 (8.8%) |  | 354 (8.9%) |
| Not tested | 19 (32.8%) | 958 (24.4%) |  | 977 (24.5%) |
| **Hepatitis co-infection, n (%)** |  |  | 0.155 |  |
| No | 30 (51.7%) | 2490 (63.5%) |  | 2520 (63.3%) |
| Yes | 8 (13.8%) | 351 (8.9%) |  | 359 (9.0%) |
| Not tested | 20 (34.5%) | 1081 (27.6%) |  | 1101 (27.7%) |
| **Calendar year of index date** |  |  | 0.476 |  |
| Median (IQR) | 2012 (2007, 2016) | 2012 (2009, 2016) |  | 2012 (2009, 2016) |
| **Age, years** |  |  | 0.005 |  |
| Median (IQR) | 41 (35, 48) | 37 (30, 44) |  | 37 (30, 44) |
| **CD4 count, cells/mm<sup>3</sup>** |  |  | <.001 |  |
| Median (IQR) | 766 (546, 1001) | 535 (384, 707) |  | 537 (385, 711) |
| **CD8 count, cells/mm<sup>3</sup>** |  |  | <.001 |  |
| Median (IQR) | 732 (499, 997) | 984 (718, 1352) |  | 978 (715, 1348) |
| **CD4/CD8 ratio** |  |  | 0.476 |  |
| Median (IQR) | 1.13 (0.72, 1.62) | 0.53 (0.36, 0.77) |  | 0.54 (0.36, 0.78) |
| **VL set point, log<sub>10</sub> copies/mL** |  |  | <.001 |  |
| Median (IQR) | 1.40 (1.30, 1.66) | 4.36 (3.79, 4.85) |  | 4.35 (3.76, 4.84) |
| **Time from HIV diagnosis to index date, months** |  |  | 0.510 |  |
| Median (IQR) | 631 (574, 677) | 635 (592, 679) |  | 634 (592, 679) |

A significantly higher prevalence of Caucasian people (p=0.002), current smokers (p=0.026) and MSM (p<0.001) was found in the viremic group. In contrast, no evidence for a difference by groups was found regarding BMI [aviremic vs viremic 24 (IQR: 22, 27) Kg/m^2^ vs 23 (IQR: 21, 25) Kg/m^2^ p=0.06], CD4/CD8 ratio [aviremic vs viremic 1.13 (IQR: 0.72, 1.66) vs 0.53 (IQR: 0.36, 0.77) p=0.476] HIV duration [aviremic vs viremic 631 (IQR: 574, 677) months vs 635 (IQR: 592, 679) months p=0.510] and hepatitic viruses serology [aviremic vs viremic HBV p: 0.427; HCV p: 0.094 and hepatitis co-infections p: 0.155].

### Unadjusted association between VLset and HDL-c, LDL-c, TC and Triglycerides plasma concentration in ART-naïve patients

Figure 2 shows the distribution of lipid values in spontaneously aviremic an viremic patients enrolled in the study. Aviremic patients showed a significantly higher level of HDL-c plasma concentration [aviremic vs viremic median 48 (IQR: 42, 62) mg/dl vs. 42 (IQR: 35, 51) mg/dl p<0.001] and total cholesterol (TC) [aviremic vs. viremic 183 (IQR: 155, 210) mg/dl vs. 166 (IQR: 142, 191) mg/dl p=0,002] compared to viremic patients. Higher LDL-c plasma concentration and lower triglicerydes (TGL) levels were found in aviremic patients compared to viremic, although the association did not reach statistical significance [aviremic vs viremic LDL-c: 111 (IQR: 87, 135) mg/dl vs. 100 (IQR: 80, 122) p= 0.087; TGL: 89 (IQR: 69, 116) mg/dl vs. 99 (IQR: 72, 142) mg/dl p=0.094]. We also evaluated the linear correlation between all lipid parameters and HIV viremia; our data shows a negative correlation between HIV viremia and HDL-c, LDL-c and TC as well as a positive correlation with TGL plasma concentrations. In particular regarding HDL-c and VLset, we observed a significant negative correlation (Pearson R^2^=0.03) and an absolute difference of -8.05 mg/dL when comparing viremic with aviremic patients (95% CI:-15.3; -0.84, p=0.03, Table 2). In contrast, there was no evidence for a difference in total cholesterol between the viremic and aviremic group from the unadjusted linear regression with total cholesterol as outcome: -14.5 mg/dl (95% CI:-38.6; +9.36), p=0.23 (Table 4).

**Table 2.** Mean HDL-C concentrations according to HIV-RNA from fitting a linear regression model.

| Models | HDL cholesterol mg/dl, Mean Difference (95% CI) |  |  |
| --- | --- | --- | --- |
|  | VLSet <=50 | VLSet >50 | per log10 VLSet higher |
| **Unadjusted** | *Ref.* | -8.05 (-15.3, -0.84) | -3.21 (-4.14, -2.27) |
| **p-value** |  | 0.029 | <.001 |
| **Adjusted<sup>1</sup>** | *Ref.* | -7.65 (-14.9, -0.44) | -3.11 (-4.06, -2.16) |
| **p-value** |  | 0.038 | <.001 |
| **Adjusted<sup>2</sup>** | *Ref.* | -5.24 (-12.4, 1.94) | -2.57 (-3.52, -1.62) |
| **p-value** |  | 0.153 | <.001 |

<sup>1</sup>for CD4/CD8, age, AIDS and HCVAbs status
<sup>2</sup>for those in model<sup>1</sup> plus gender

**Table 3.**
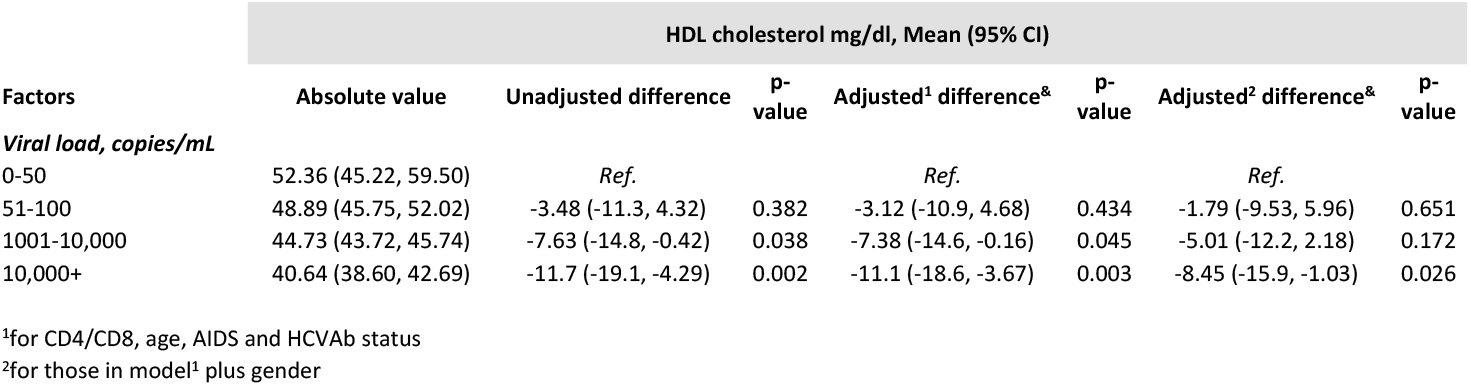
Mean HDL-C concentrations according to HIV-RNA from fitting a linear regression model dose-response model.

| Factors | HDL cholesterol mg/dl, Mean (95% CI) |  |  |  |  |  |  |
| --- | --- | --- | --- | --- | --- | --- | --- |
|  | Absolute value | Unadjusted difference | p-value | Adjusted<sup>1</sup> difference<sup>&</sup> | p-value | Adjusted<sup>2</sup> difference<sup>&</sup> | p-value |
| **Viral load, copies/mL** |  |  |  |  |  |  |  |
| 0-50 | 52.36 (45.22, 59.50) | *Ref.* |  | *Ref.* |  | *Ref.* |  |
| 51-100 | 48.89 (45.75, 52.02) | -3.48 (-11.3, 4.32) | 0.382 | -3.12 (-10.9, 4.68) | 0.434 | -1.79 (-9.53, 5.96) | 0.651 |
| 1001-10,000 | 44.73 (43.72, 45.74) | -7.63 (-14.8, -0.42) | 0.038 | -7.38 (-14.6, -0.16) | 0.045 | -5.01 (-12.2, 2.18) | 0.172 |
| 10,000+ | 40.64 (38.60, 42.69) | -11.7 (-19.1, -4.29) | 0.002 | -11.1 (-18.6, -3.67) | 0.003 | -8.45 (-15.9, -1.03) | 0.026 |

<sup>1</sup>for CD4/CD8, age, AIDS and HCVAbs status
<sup>2</sup>for those in model<sup>1</sup> plus gender

**Table 4.** Mean total Cholesterol concentrations according to HIV-RNA from fitting a linear regression model.

| Models | HDL cholesterol mg/dl, Mean Difference (95% CI) |  |  |
| --- | --- | --- | --- |
|  | VLSet <=50 | VLSet >50 | per log10 VLSet higher |
| **Unadjusted** | *Ref.* | -14.6 (-38.6, 9.36) | -4.46 (-7.59, -1.33) |
| **p-value** |  | 0.232 | 0.005 |
| **Adjusted<sup>1</sup>** | *Ref.* | -11.2 (-35.1, 12.67) | -4.36 (-7.52, -1.20) |
| **p-value** |  | 0.357 | 0.007 |
| **Adjusted<sup>2</sup>** | *Ref.* | -8.96 (-32.9, 15.00) | -3.88 (-7.07, -0.69) |
| **p-value** |  | 0.464 | 0.017 |

<sup>1</sup>for CD4/CD8, age, AIDS and HCVAbs status
<sup>2</sup>for those in model<sup>1</sup> plus gender

**Figure 2.**
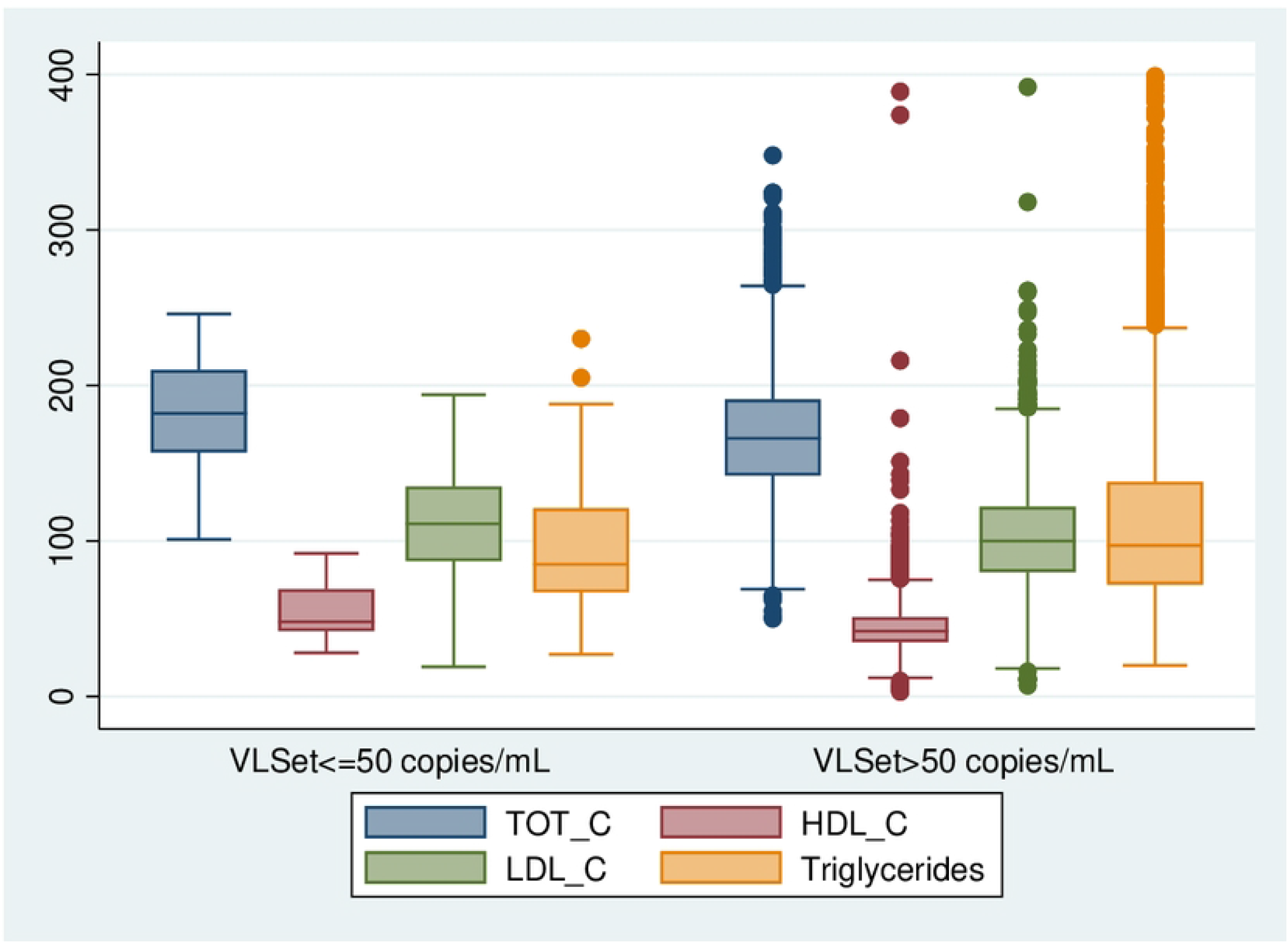
Box plots of biomarkers according to Vlset.

### Role of potential confounding factors

The relationship between VLset and HDL-c and TC was re-evaluated after controlling for potential confounders using linear regression adjustment. When VLSet was fitted as a binary exposure (aviremic vs. viremic) it was associated with HDL-C levels independently of age, AIDS diagnosis and HCVAb status. However, after controlling for gender this effect was somewhat attenuated (Table 2). This is because females are known to have a lower VLSet [26] and also a higher HDL-C. Interestingly, confounding was less strong in the analysis in which VLSet was fitted as continuous in the log_10_ scale which has greater statistical power. Also, difference could still be seen when comparing aviremic patients with those with very high levels of HIV-RNA (>10,000 copies/mL), even after controlling for gender (Table 3). Table 3 also shows a nice dose-response relationship between HIV-RNA and HDL-c which, despite the cross-sectional nature of the analysis, seems to suggest causality. In the main analysis with VLSet fitted as continuous in the log scale, with an observed standardised difference of 2.57 logs in the fully adjusted model and a standard error for this difference of 0.49, an unmeasured confounder that was associated with both the outcome and the exposure each with a log difference of at least 20.2 logs could explain away the estimate, but weaker confounding could not. Similarly, to move the confidence interval to include the null, an unmeasured confounder that was associated with the outcome and the exposure each by a difference of at least 8.1 logs could do so, but weaker confounding could not. To put this in prospective, the difference associated with the measured factors showing the strongest association was 9.3 logs for gender.

In contrast, the model with TC as outcome showed an association with VLSet only when the latter was fitted as continuous in the log_10_ scale (Table 4). The analysis show that other factors such as age, AIDS and HCVAb status played a role in explaining the unadjusted difference in total cholesterol between the aviremic and viremic group.

### Mediation analysis

We further evaluated the total direct effect of VLSet on CD4/CD8 ratio by decomposing the effect in the direct effect of VLSet on CD4/CD8 ratio and the indirect effect through the causal pathway of HDL-C (Figure 1). This analysis indicated that indeed some of the total effect of VLSet on CD4/CD8 is significantly mediated by a variation in HDL-c induced by HIV-RNA. Although significant, this indirect effect is estimated to be only a small percentage of the total effect (Table 5). There was also evidence that the indirect effect was larger, although still small in absolute terms, in people with lower levels of HDL-c which was estimated after formally testing for interaction (data not shown).

**Tab 5.** Results of the mediation analysis with outcome CD4/CD8.

**Tab 5 Results of the mediation analysis with outcome CD4/CD8**
| **VL Set-point (per log10 higher)** |  |  |  |
| --- | --- | --- | --- |
|  | **Mean CD4/CD8** | **95% CI** | **p-value** |
| **Direct Effect** | -0.13 | -0.14; -0.11 | <0.0001 |
| **Indirect effect via HDL-C** | -0.0009 | -0.002; -0.0003 |  |
| **Total Effect** | -0.13 | -0.14; -0.11 |  |
| **% of Tot Effect mediated by HDL-C** | 0.69% | 0.61%; 0.79% |  |

Mediation analysis on the assumptions described in the DAG (Figure 1).

## Discussion

In our retrospective cross-sectional analysis, for the first time on a large sample of real-life untreated PLWH, we found evidence for a significant inverse relationship in vivo between HDL-c plasma concentration and HIV viremia.

Regarding the important role that lipoproteins assume in infectious diseases, there is indeed evidence for a strict relationship between lipid metabolism and viral replication. Specifically, membrane cholesterol-rich lipid rafts have multiple functions for viral replication, recruiting and concentrate several receptors and molecules involved in pathogen recognition and cellular signalling, which mediate pathogen internalization and modulate the lipid raft-dependent immune response.

Focusing on the results of our analysis, we found an inverse relationship between HDL-c levels and VLSet which could not be fully explained by a number of key measured confounding variables. Higher levels of HIV-RNA were associated with a lower HDL-c independently of age, CD4/CD8 ratio, AIDS and HCVAb status; despite the cross-sectional nature of the study design, under our strong assumptions of a correctly specified model and no unmeasured confounding, the observed link could be interpreted as causal in that low levels of HDL-c are determined by higher levels of HIV replication. When grouping study participants as viremic vs. aviremic, the difference was largely attenuated by gender, although this analysis is likely to have low statistical power. No association was detected between VLset and LDL-c, while a significant association between VLset and both TGL and TC in unadjusted analysis was largely explained by confounding factors. Especially when dealing with observational data it is important to question whether the findings might be due to bias and these other results, which act as negative controls, are somewhat in support of the evidence. Overall, our results appear to confirm the presence of a link between HIV replication and lipid metabolism. In particular, we speculate that the inverse correlation seen between HIV viremia and HDL-c in our “in vivo” study, is a result of the fact Nef HIV protein was able, through active viral replication, to reduce HDL-c production by impairing ABCA-1, generating cholesterol accumulation within macrophages, promoting their foam-cells transformation and increasing the cardiovascular risk among PLWH (27, 28).

Moreover, HIV-RNA is known to have a direct effect on immune-parameters such as CD4 count, CD8 and their ratio [29-33]. On the basis of the results of our analysis, we could also speculate that higher HIV-RNA replication may cause a reduction of HDL-c levels, which in turn leads to higher level of inflammation markers (e.g. cytokines and monocyte activation), with a further effect in reducing the CD4/CD8 ratio. Our formal mediation analysis supports the existence of this indirect effect although it represents only a very small percentage of the total effect. In general, this result reinforces our hypothesis of the role of HIV-replication in causing lipids and immunological abnormalities.

It is known that HDL-c might decrease the expression of several key components of the inflammasomes during HIV infection, suggesting a crucial role of HDL-c in modulate the inflammatory state and consequently, the progression of HIV infection. Moreover, greater interleukin-6 (IL-6) and intercellular adhesion molecule-1 (ICAM-1) levels have been recently found to be associated with both lower total HDL-c and small HDL particles.[27-28]. Further studies are needed to better evaluate the association between lower HDL-c and small HDL particles on IL-6 and other cytokines (which were not included in our analysis because they were available only for the subset of the aviremic individuals), considering also the potential contribute of these mechanisms to increased CVD risk among PLWH [34-36].

Reasons for the increased risk in CVD in PLWH as compared to that observed in the general population remain still partly unclear. Our data suggest that HIV replication alone could have a pivotal role in increasing this risk by its direct effect on HDL-c reduction and triglycerides elevation, independently of ART. Other studies should be conceived to further evaluate the causal link between HIV-RNA and the risk of CVD, carefully investigating the role of HIV-RNA as the main exposure of interest, ART and HCV-RNA as key confounding factors, and HDL-c as the potential key mediator; in contrast, most analysis thus far have considered lipids elevation, perhaps wrongly, as a confounder for the effect of ART instead of being a mediator.

Before drawing final conclusions, a number of limitations of our analysis need to be mentioned. First, although HDL particles play a critical role in the maintenance of cholesterol balance in the arterial wall and in reduction of pro-inflammatory responses by arterial cholesterol loaded macrophages, their plasmatic concentration is not a perfect surrogate marker for macrophage cholesterol efflux. Therefore, it is possible that HDL-c as routinely measured in the clinics is not a perfect surrogate of cellular cholesterol efflux and measurement error for the outcome in our analysis might exist. However, this is potentially a conservative bias as it implies that the magnitude of the association could have been diluted.

In addition, our analysis of the possible causal effect of VLSet on HDL-C is based on the assumption of no unmeasured confounding and correct specification of our model (e.g. one of the underlying assumption of our model is that BMI is a predictor of outcome but not a cause of variation in VLSet, etc. see Figure 1). However unmeasured confounding can never be ruled out in real-world data. For example, HCV-RNA at ART initiation which is not available in the database for the majority of our participants is a potential key unmeasured confounding factor. Nevertheless, many important measured confounders have been accounted for and our sensitivity analysis (through calculation of the e-value) shows that results are fairly robust to potential unmeasured confounding bias. Similar considerations apply also to the second part of our analysis, aiming to estimate the indirect and direct effect of VLSet on CD4/CD8 ratio and even more so as one key assumption in mediation analysis is that there is no mediator-outcome unmeasured confounding. Furthermore, there are many different factors that could influence our main exposure/intervention variable (individuals’ HIV-RNA set-point levels) and, in this situation according to some, one of the key conditions for the identifiability of causal effects from observational data does not hold [37]. More in general, given the cross-sectional design of the study, it is impossible to establish the exact temporality between VLSet and HDL-c and it is an arbitrary assumption, based on the exact dates of biomarkers, that we have modelled HDL-c (outcome) as a function of VLSet (exposure) and not viceversa.

In conclusion, our data show that HDL-c plasma concentration is significantly lower in absence of ART in viremic compared to aviremic patients, although this association was in part explained by gender. Further analyses should be promoted in order to study the molecular mechanisms that underline the relationship between HIV replication, HDL-c formation, and diseases progression and the role of HIV replication alone in increasing the risk of CVD in the HIV-infected population.

## Icona Foundation Study Group

### BOARD OF DIRECTORS

A d’Arminio Monforte (President), A Antinori (Vice-President), S Antinori, A Castagna, R Cauda, G Di Perri, E Girardi, R Iardino, A Lazzarin, GC Marchetti, C Mussini, E Quiros-Roldan, L Sarmati, B Suligoi, F von Schloesser, P Viale.

### SCIENTIFIC SECRETARY

A d’Arminio Monforte, A Antinori, A Castagna, F Ceccherini-Silberstein, A Cingolani, A Cozzi-Lepri, E Girardi, A Gori, S Lo Caputo, G Marchetti, F Maggiolo, C Mussini, M Puoti, CF Perno.

### STEERING COMMITTEE

C Agrati, A Antinori, F Bai, A Bandera, S Bonora, A Calcagno, D Canetti, A Castagna, F Ceccherini-Silberstein, A Cervo, S Cicalini, A Cingolani, P Cinque, A Cozzi-Lepri, A d’Arminio Monforte, A Di Biagio, R Gagliardini, A Giacomelli, E Girardi, N Gianotti, A Gori, G Guaraldi, S Lanini, G Lapadula, M Lichtner, A Lai, S Lo Caputo, G Madeddu, F Maggiolo, V Malagnino, G Marchetti, C Mussini, S Nozza, CF Perno, S Piconi, C Pinnetti, M Puoti, E Quiros Roldan, R Rossotti, S Rusconi, MM Santoro, A Saracino, L Sarmati, V Spagnuolo, N Squillace, V Svicher, L Taramasso, A Vergori.

### STATISTICAL AND MONITORING TEAM

F Bovis, A Cozzi-Lepri, I Fanti, A Rodano’, M Ponzano, A Tavelli.

### COMMUNITY ADVISORY BOARD

A Bove, M Cernuschi, L Cosmaro, M Errico, A Perziano, V Calvino.

### BIOLOGICAL BANK INMI AND SAN PAOLO

S Carrara, S Graziano, G Prota, S Truffa, D Vincenti, Y D’Errico.

### PARTICIPATING PHYSICIANS AND CENTERS

Italy A Giacometti, A Costantini, V Barocci (Ancona); A Saracino, C Santoro, E Milano (Bari); F Maggiolo, C Suardi (Bergamo); P Viale, L Badia, S Cretella (Bologna); E Quiros Roldan, E Focà, C Minardi (Brescia); B Menzaghi, C Abeli (Busto Arsizio); L Chessa, F Pes (Cagliari); P Maggi, L Alessio (Caserta); B Cacopardo, B Celesia (Catania); J Vecchiet, K Falasca (Chieti); A Pan, S Dal Zoppo (Cremona); D Segala (Ferrara); F Vichi, MA Di Pietro (Firenze); T Santantonio, S Ferrara (Foggia); M Bassetti, E Pontali, S Blanchi, N Bobbio, G Mazzarello (Genova); M Lichtner, L Fondaco (Latina); S Piconi, C Molteni (Lecco); S Rusconi, G Canavesi (Legnano) A Chiodera, P Milini (Macerata); G Nunnari, G Pellicanò (Messina); A d’Arminio Monforte, S Antinori, A Lazzarin, G Rizzardini, M Puoti, A Gori, A Castagna, A Bandera, V Bono, MV Cossu, A Giacomelli, R Lolatto, MC Moioli, L Pezzati, C Tincati (Milano); C Mussini, C Puzzolante (Modena); P Bonfanti, G Lapadula (Monza); V Sangiovanni, I Gentile, V Esposito, FM Fusco, G Di Filippo, V Rizzo, N Sangiovanni (Napoli); AM Cattelan, S Marinello (Padova); A Cascio, C Colomba (Palermo); D Francisci, E Schiaroli (Perugia); G Parruti, F Sozio (Pescara); P Blanc, A Vivarelli (Pistoia); C Lazzaretti, R Corsini (Reggio Emilia); M Andreoni, A Antinori, R Cauda, C Mastroianni, A Cingolani, V Mazzotta, S Lamonica, M Capozzi, A Mondi, M Rivano Capparuccia, G Iaiani, C Stingone, L Gianserra, J Paulicelli, MM Plazzi, G d’Ettore, M Fusto (Roma); M Cecchetto, F Viviani (Rovigo); G Madeddu, A De Vito (Sassari); M Fabbiani, F Montagnani (Siena); A Franco, R Fontana Del Vecchio (Siracusa); BM Pasticci, C Di Giuli (Terni); GC Orofino, G Calleri, G Di Perri, S Bonora, G Accardo (Torino); C Tascini, A Londero (Udine); V Manfrin, G Battagin (Vicenza); G Starnini, D Farinacci (Viterbo).

